# Improved Statistical Efficiency of Simultaneous Multi-Slice fMRI by Reconstruction with Spatially Adaptive Temporal Smoothing

**DOI:** 10.1101/646554

**Authors:** Mark Chiew, Karla L. Miller

## Abstract

We introduce an approach to reconstruction of simultaneous multi-slice (SMS)-fMRI data that improves statistical efficiency. The method incorporates regularization to adjust temporal smoothness in a spatially varying, encoding-dependent manner, reducing the g-factor noise amplification per temporal degree of freedom. This results in a net improvement in tSNR and GLM efficiency, where the efficiency gain can be derived analytically as a function of the encoding and reconstruction parameters. Residual slice leakage and aliasing is limited when fMRI signal energy is dominated by low frequencies. Analytical predictions, simulated and experimental results demonstrate a marked improvement in statistical efficiency in the temporally regularized reconstructions compared to conventional slice-GRAPPA reconstructions, particularly in central brain regions. Furthermore, experimental results confirm that residual slice leakage and aliasing errors are not noticeably increased compared to slice-GRAPPA reconstruction. This approach to temporally regularized image reconstruction in SMS-fMRI improves statistical power, and allows for explicit choice of reconstruction parameters by directly assessing their impact on noise variance per degree of freedom.

## 1. Introduction

Simultaneous multi-slice (SMS) imaging (Barth et al., 2016; Larkman et al., 2001; Moeller et al., 2010) has become widely used for functional MRI (fMRI) data acquisition, particularly since the introduction of the controlled aliasing in parallel imaging (CAIPI) (Breuer et al., 2005) blipped EPI acquisition (Setsompop et al., 2012). The blipped CAIPI scheme significantly reduced noise amplification g-factor penalties associated with the SMS unaliasing by introducing a 3D k-space sampling pattern (Zahneisen et al., 2014; Zhu et al., 2012) that manipulates aliasing locations to improve conditioning of the encoding matrix.

All SMS reconstruction methods rely on knowledge of the coil sensitivities (Pruessmann et al., 1999) or equivalently, the GRAPPA kernels (Griswold et al., 2002) for reconstruction. Broadly speaking, SMS reconstructions have employed either SENSE/GRAPPA (Blaimer et al., 2013, 2006; Koopmans, 2017; Moeller et al., 2014), Slice-GRAPPA (Cauley et al., 2014; Hoge et al., 2018; Setsompop et al., 2012), or conventional SENSE (Zahneisen et al., 2014) formulations. In addition to coil sensitivity information, SMS reconstruction using additional phase (Blaimer et al., 2013) or low-rank (coil x space) (Kim and Haldar, 2015; Park and Park, 2017) constraints have also been proposed. Some of these methods have been specifically designed to minimize reconstruction spatial bias (Cauley et al., 2014; Park and Park, 2017) that arises from inter-slice leakage artifacts.

All of the above methods operate in a time-independent manner, reconstructing images individually. In the context of functional MRI (fMRI), methods additionally exploiting temporal structure in the data have been demonstrated in compressed sensing approaches, (Chavarrías et al., 2015; Jung et al., 2007; Zong et al., 2014), or by minimizing image differences (Li et al., 2018). Alternatively, low-rank (space x time) methods have been proposed for use in fMRI based on the use of low-dimensionality representations in fMRI analysis methods (Chiew et al., 2016, 2015). While low-rank approaches do not impose any specific constraint on the representation of spatial or temporal information, more recent low-rank plus sparse methods have also been used to additionally exploit sparsity in the temporal Fourier domain (Aggarwal et al., 2017; Petrov et al., 2017; Singh et al., 2015; Weizman et al., 2017) which requires smoothness or periodicity in voxel time-courses.

Here, we introduce an improvement to SMS-EPI for fMRI by introducing time-varying sampling with a spatially adaptive temporally regularized reconstruction. Analytical expressions for reconstruction bias and variance are presented, as well as assessment of the relative statistical efficiencies of a general linear model (GLM) analysis, and temporal SNR efficiency as a function of the image reconstruction parameters. We exploit the low-frequency distribution of BOLD signals to ensure that the proposed reconstruction results in comparable or lower mean squared error than is produced by conventional SENSE or slice-GRAPPA unaliasing, while improving statistical power. This paper demonstrates the utility of this proposed reconstruction in simulations and experiments, and provides a framework for principled analysis of temporal regularization factors in fMRI by directly calculating the impact of unaliasing reconstruction parameters on GLM analysis.

## 2. Theory

### 2.1 Linear Reconstruction

By considering SMS unaliasing to be a special case of a 3D under-sampling problem, a general SENSE-based framework can be used for reconstruction. The linear reconstruction problem we propose to solve is:

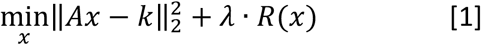

where *A* is the linear encoding operator, which incorporates point-wise multiplication of coil sensitivities (assumed to be whitened) and the k-space sampling transform (which can be Cartesian or non-uniformly sampled). If the spatio-temporal image vector *x* is formed by vertically concatenating the time-series images across time-points (into a *N_x_N_y_N_z_N_t_* × 1 vector, with *N_x,y,z_* corresponding to spatial dimensions *x, y* and *z*, and *N_t_* corresponding to the number of time-points), then *A* is a block-diagonal matrix where the block corresponding to time-point *t* is 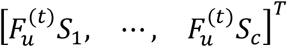, with 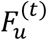 denoting the under-sampled Fourier transform at time *t*, and *S_D_* is the sensitivity for the *i^th^* coil, for 1 < *i* < *c*, where *c* is the number of coils. In Eq. 1, *k* is the vectorized noisy k-space data, *R* is the L2 penalty that enforces temporal smoothing, and *λ* is the parameter that controls the amount of smoothing and effective temporal resolution.

We choose *R* to be:

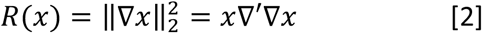

where ∇ is the finite difference operator acting on the time domain (voxel-by-voxel), and ∇’ denotes the adjoint of ∇. The L2 penalty allows the consequence of *R* to be interpreted like a linear smoothing operation, with the solution to Eq. 1 well known to be:

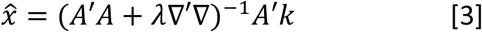

### 2.2 Reconstruction Bias and Variance

To characterize the estimator in Eq. 3, we consider the total mean squared reconstruction error as *MSE = bias*^2^ + *variance*, where the bias term represents signal misallocation error (e.g. leakage or residual aliasing) and the variance term related to noise amplification (e.g. g-factor). We first reformulate Eq. 3 in terms of an equivalent, encoding dependent smoothing operator, *S_λ_*, that is applied following a conventional pseudoinverse reconstruction:

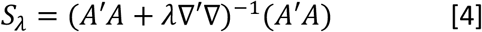

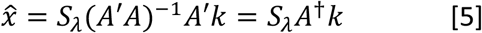

where *A*^†^ denotes the Moore-Penrose pseudoinverse of *A*. Using this formulation, it is apparent that *S_λ_* is spatially adaptive in that it depends on *A’A* (Fessler and Rogers, 1996), unlike conventional shift-invariant post-hoc kernel smoothing.

With a conventional unbiased pseudoinverse reconstruction, *E*(*A*^†^*k*) = *E*(*x*) = *x*, and so the bias expected from this reconstruction can be described by:

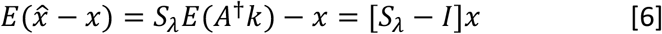

The g-factor at each time-point can be derived from the square root of the voxel-wise ratio of the variance of the estimator 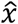 and the variance of a fully sampled, unsmoothed reconstruction (see Eq. A9, Appendix A):

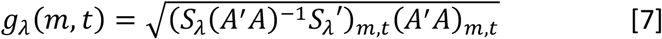

where here *M_m,t_* denotes the diagonal element of any matrix *M* corresponding to the *m^th^* voxel and *t^th^* time-point, resulting in a g-factor for every voxel and time-point. The average g-factor for any voxel *m* is then given by the root-mean-square across time (Ramb et al., 2015):

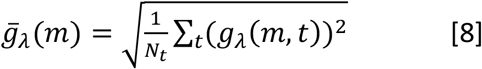

### 2.3 GLM Efficiency

The benefit of regularization, in the context of general linear model regression analysis, can be quantified by the efficiency of the estimator produced from the proposed reconstruction. Efficiency is inversely related to the variance of the estimated GLM regression coefficient, and an increase in efficiency results in increased statistical power.

A complete derivation of the relative efficiency between a regularized and un-regularized reconstruction can be found in Appendix B, following closely from Worsley and Friston (Worsley and Friston, 1995). Efficiency is a voxel-wise measure depending on g-factor, *S_λ_*, and the task design matrix:

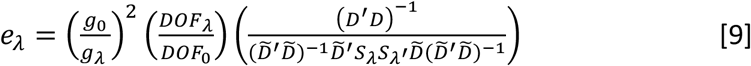

where

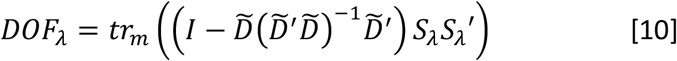

can be interpreted as an effective degrees of freedom (DOF) measure, where 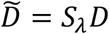 is the smoothed task design matrix *D*, and *tr_m_*(·) indicates that values across time for a given voxel *m* are summed.

Asymptotically, when *N_t_* is sufficiently large, *λ* is small, and the task waveform is sufficiently slowly varying, Eq. 9 simplifies to an expression independent of *D*:

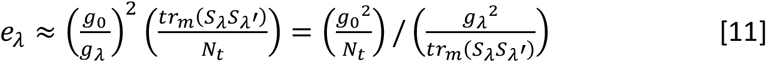

With *λ* = 0 (no smoothing), *tr_m_*(*S_λ_S_λ_*′) = *N_t_* for all voxels, which naturally leads to defining an approximate DOF expression as:

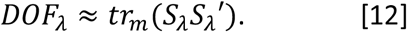

Then Eq. 11 can be interpreted intuitively as the ratio of the squared g-factors (*g*^2^) normalized by their respective DOF. When regularization reduces g-factor faster than the loss of effective DOF, efficiency and statistical power are increased compared to the unregularized case.

### 2.4 tSNR Efficiency

Empirical temporal SNR (tSNR) efficiency can also be computed from resting condition data by adjusting the voxel-wise temporal standard deviation with the normalized DOF using the approximate form in Eq. 12:

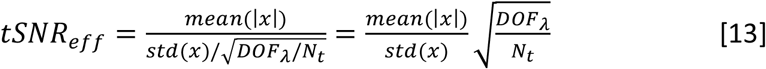

with the mean and standard deviation (*std*) taken across time. When *DOF_λ_*/*N_t_* = 1, this is equivalent to conventional tSNR definitions, but when *DOF_λ_*/*N_t_* < 1, tSNR efficiency decreases to reflect the reduced degrees of freedom per unit time.

## 3. Methods

To characterize the proposed method, numerical simulations and experiments were performed to assess the performance of the proposed reconstruction method.

First, two sets of numerical simulations were used to separately characterize noise and signal properties, exploiting the linearity of the reconstruction approach:

***Sim1.*** A “noise only” Monte Carlo simulation to estimate the spectral response to regularization and validate the analytical g-factor, DOF and efficiency expressions
***Sim2.*** A “signal only” numerical simulation to assess signal bias or leakage in the absence of any noise amplification effects

Next, three experiments were performed on healthy volunteers to assess the reconstruction performance *in vivo*, using different acquisition protocols:

***Exp1.*** Resting fMRI with single-slice excitation and full SMS reconstruction, which allows for the evaluation of total mean squared error in the unexcited slices
***Exp2.*** Resting fMRI with SMS-excitation and reconstruction, to evaluate tSNR efficiency
***Exp3.*** Task fMRI with SMS-excitation and reconstruction, to evaluate GLM statistical efficiency
***Exp4.*** Resting fMRI with SMS-excitation and reconstruction, to evaluate tSNR efficiency in thinner slices, comparing protocols with fixed TRvol

### 3.1 CAIPI Sampling

The k-space sampling schemes explored in this work can be characterized by 3 parameters:

Δ*kz*/Δ*ky*: defines the conventional *within-shot* CAIPI sampling pattern, corresponding to the z-gradient blips that control the FOV shift between simultaneously acquired slices
Δ*kz*/Δ*t*: defines the *between-shot* shift of the sampling pattern in the *kz* direction, which corresponds to a changing phase relationship between simultaneously acquired slices
Δ*ky*/Δ*t*: defines the *between-shot* shift of the sampling pattern in the *ky* direction, which is only non-zero if in-plane acceleration is used

Figure 1 illustrates the effects of Δ*kz*/Δ*ky* = 3/2, Δ*kz*/Δ*t* = 3, and Δ*ky*/Δ*t* = 0 sampling pattern. For SMS acquisitions with conventional CAIPI sampling, Δ*kz*/Δ*t* = 0, which means the sampling pattern is identical from shot-to-shot. Here, by introducing sampling patterns where Δ*kz*/Δ*t* and Δ*ky*/Δ*t* are non-zero, the sampling pattern (defined by Δ*kz*/Δ*ky*) is shifted in k-space from shot-to-shot. Although these parameters can also vary with time, in this work, we focus mainly on acquisitions with constant sampling parameters, so that the relative shift from any shot to the next does not change.

**Figure 1.**
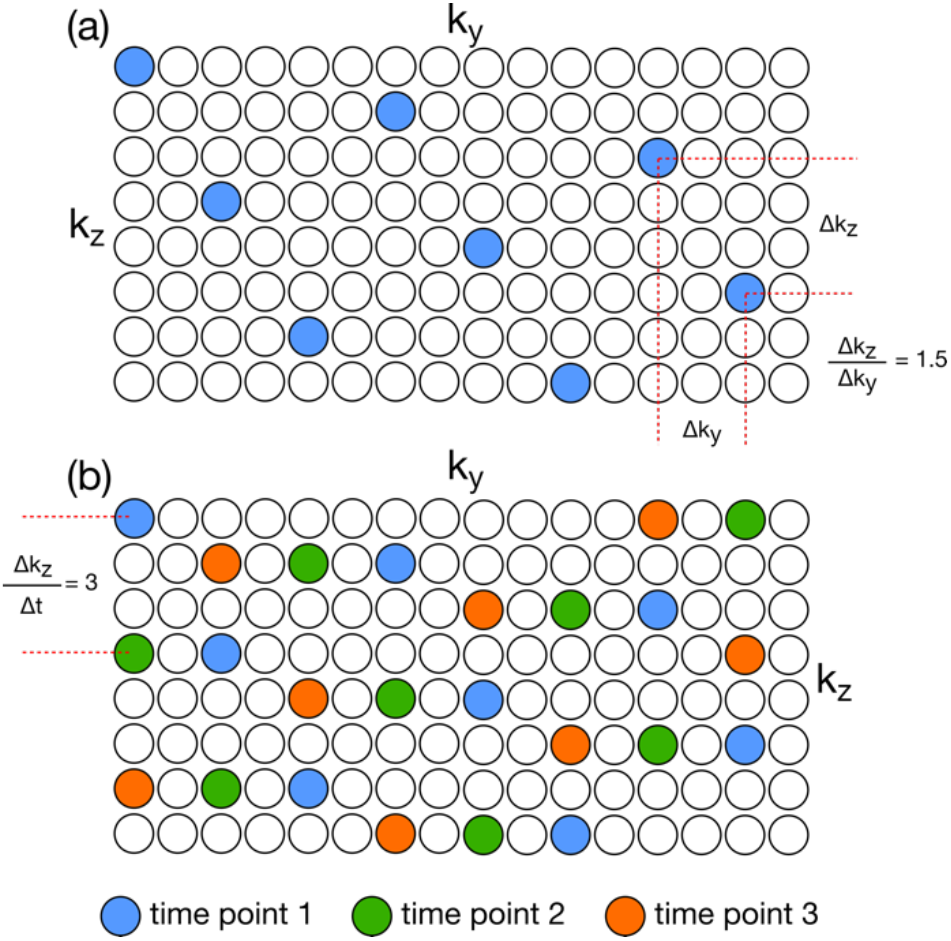
(a) Example CAIPI sampling pattern for a MB8R2 acquisition. In this example, the Δk_z_ = 3, and Δk_y_ = 2, leading to a slice-wise CAIPI shift of FOV*3/16. (b) The sampling pattern can be shifted over time, as shown here by a Δk_z_/Δt = 3. The time-shifting parameter Δk_z_/Δt does not need to be constant, and can be pseudorandom.

One advantage of designing the sampling scheme with constant shift parameters (i.e. regular shifts across time) is that this results in g-factors that are also constant across time, simplifying analysis and ensuring a stationary noise model for the entire time-series reconstruction. Intuitively, this results from the property that any width sampling window centered on any given time-point will have sampling patterns that are identical, to within a shift, and that g-factors are invariant to shifts in k-space.

### 3.2 Numerical Simulations

To form a basis for both sets of numerical simulations, a single-band reference dataset was acquired on a cylindrical phantom using a 64-channel head and neck coil. This reference data was used to provide realistic coil sensitivities and noise covariance, and to define the geometry of the image for the signal modelled in *Sim2*. Both *Sim1* and *Sim2* simulated the same MB=8 acquisition with no in-plane acceleration, and 256 timepoints using 16 compressed virtual coils using the geometric coil compression scheme (Zhang et al., 2013).

#### 3.2.1 Simulation 1 (*Sim1*)

In *Sim1*, the Monte Carlo simulations were performed using Gaussian white additive noise across 1000 different complex noise realizations. To compare the proposed reconstructions against post-hoc smoothing, linear smoothing operators *S_k_* = (*I* + *k*∇’∇)^−1^ were constructed, with *κ* chosen empirically to match effective DOF with *S_λ_* using Eq. 10, across a range of *λ* spanning 10^−5^ to 10^2^, and (Δkz/Δ*ky* = 3, Δkz/Δ*t* = 2) sampling. Three different GLM design regressors were initially used to evaluate DOF and efficiency: (i) 5-period block design, (ii) fast event-related design, (iii) white noise regressor (see Supporting Figure S1).

#### 3.2.1 Simulation 2 (*Sim2*)

In *Sim2*, inter-slice leakage was examined by simulating reconstruction of a noise-free dataset with a voxel-wise random signal, repeated 10 times. A 1/f characteristic of BOLD fMRI data (Zarahn et al., 1997) was used to provide a low-frequency signal model for assessing leakage bias, because residual bias is signal-dependent (Eq. 6) in the proposed reconstruction. To characterize leakage, in this simulation only slice 5 contained signal, such that any reconstructed signal in the other 7 slices would have to come from signal leakage. A range of different *λ* (from 0 to 10^−2^) and sampling patterns (∇k*z*/∇*ky* = 3, ∇k*z*/∇*t* = 0,1, 2, 3) were explored, with efficiencies calculated using the event-related regressor only.

### 3.3 In Vivo Experiments

Four total subjects were scanned on a 3 T system (Prisma, Siemens Healthineers) using a 64-channel head and neck coil with informed consent in accordance with local ethics. Acquisition parameters common to experiments 1-3 were: 2 mm isotropic resolution, 64 slices, flip angle = 40°, phase encoding direction = AP, bandwidth = 2368 Hz/px. For the “MB8R1” protocols (Δ*kz*/Δ*ky* = 5 and Δ*kz*/Δ*t* = 2), sets of 8 slices were acquired simultaneously, with no in-plane acceleration, with TE = 38 ms and TRvol = 680 ms. For the “MB8R2” protocols (Δ*kz*/Δ*ky* = 5/2, Δ*kz*/Δ*t* = 3, Δ*ky*/Δ*t* = 1), sets of 8 slices were acquired simultaneously, with R=2 in-plane, with TE = 30 ms and TRvol = 520 ms. MB excitation pulses were designed to minimize peak power using optimized phase schedules (Wong, 2012). Sensitivity maps were estimated using the ESPIRiT method (Uecker et al., 2014), from a separate single-band reference acquisition, and the same reference data was used to train slice GRAPPA kernels. All data were compressed down to 24 virtual channels after whitening with the coil noise covariance.

#### 3.3.1 Experiment 1 (Exp1)

In *Exp1*, one subject was scanned using the MB8R1 and MB8R2 protocols in a resting condition, except the MB excitations were replaced by single slice excitations, of slice 5 only. This data was reconstructed using the full SMS reconstruction to recover 8 imaging slices (without using any knowledge that the other 7 slices are unexcited). This allowed for an assessment of total mean squared error (bias^2^ + noise variance) in the unexcited slices. Voxel-wise assessment of efficiency gains in the excited slice were also calculated, using a reference reconstruction generated with the knowledge that only a single slice has actually been excited, and performing a standard, non-SMS reconstruction of the same data (without any noise amplification due to the SMS unaliasing). One minute of data for each (96 and 128 time-points for MB8R1 and MB8R2 respectively) were acquired.

#### 3.3.2 Experiment 2 (*Exp2*)

In *Exp2*, one subject was scanned using the same MB8R1 and MB8R2 protocols as in *Exp1*, but with full MB=8 excitations. One minute of data for each (96 and 128 time-points for MB8R1 and MB8R2 respectively) were acquired, which was used to estimate voxel-wise maps of resting tSNR efficiency.

#### 3.3.3 Experiment 3 (*Exp3*)

In *Exp3*, two subjects were scanned in a 5 minute, 30 s off/on block design visual task experiment. One subject using the MB8R1 protocol (440 time-points), and a second subject using the MB8R2 protocol (576 time-points). The functional data were analyzed using FSL FEAT, with pre-whitening for noise auto-correlations (Woolrich et al., 2001).

#### 3.3.4 Experiment 4 (*Exp4*)

In *Exp4*, to evaluate reconstruction performance in fixed-TR (and fixed temporal autocorrelation) conditions, one subject was scanned in two whole-brain, one-minute resting state acquisitions to assess tSNR efficiency. A “MB8 – 50% gap” protocol with 8 sets of 8 simultaneous (64 total) slices, with Δ*kz*/Δ*ky* = 5 and Δ*kz*/Δ*t* = 2, and a “MB12 – no gap” protocol with 8 sets of 12 simultaneous (96 total) slices, with Δ*kz*/Δ*ky* = 5 and Δ*kz*/Δ*t* = 3. The acquisitions used a slice thickness of 1.5 mm, which reduces through-plane dephasing and increases the thermal noise dominance. In both cases, coverage in the superior-inferior direction extended 144 mm, but in the “MB8 – 50% gap” acquisition, a slice gap of 50% (.75 mm) was used to maintain a fixed TRvol = 712 ms. Additional parameters for these acquisitions were: TE = 39 ms (no in-plane acceleration) and bandwidth = 2480 Hz/px.

### 3.4 SMS-Reconstruction

Eq. 4 was solved iteratively using a conjugate gradient algorithm implemented in MATLAB, using a tolerance of 10^−4^ and a maximum number of iterations of 200. Reconstruction times depend on the data and *λ*, but typically ranged between 30 minutes – 2 hours for a 96×96×8×256 slice group on a 32-core AMD Opteron computer. The g-factor and efficiency expressions were evaluated directly, by partitioning the matrix inversions into smaller independent subproblems by exploiting the independence of each x-location (along the fully-sampled readout direction) and the coherent aliasing patterns when possible. Reconstruction code for the proposed method can be found at https://users.fmrib.ox.ac.uk/~mchiew/research. The *in vivo* data were compared with conventional slice-GRAPPA reconstructions using a MATLAB implementation of the CMRR-SMS (Center for Magnetic Resonance Research, University of Minnesota) reconstruction with the LeakBlock option set. All data were magnitude transformed following image reconstruction, which constitutes a non-linear operation on the data and alters the noise distribution, but this effect should be negligible when tSNR > 2 (Gudbjartsson and Patz, 1995).

## 4. Results

The results of *Sim1* are plotted in Figure 2. In Fig. 2a, noise spectra are plotted, normalized to the unregularized spectrum, for *λ* = 0, 1 × 10^−3^ and 1 × 10^−2^ reconstructions, and DOF-matching post-hoc smoothing *κ* = (2.89 × 10^−2^,1.66 × 10^−1^). As total noise is the sum of squares in either the temporal or frequency domain, it is apparent that the regularized reconstructions result in less noise compared to post-hoc smoothing with the same effective DOF (blue and black lines having similar spectral profiles, but with the blue lines having lower amplitude than the corresponding black lines). As *λ* or *κ* increases, the effective DOF decreases and the coloring of the noise spectrum is more pronounced as higher frequency content is suppressed. Additional plots of reconstruction noise spectra as a function of *λ* can be found in Supporting Figure S2. Figs. 2b and 2c plot the variation in approximate DOF (Eq. 12) and g-factor (Eq. 8) respectively against *λ*, showing the expected decrease with increasing regularization across the three voxels (selected to reflect a range of behavior). Voxels 1 and 2 show high and moderate g-factor voxels respectively, while voxel 3 does not alias onto any other voxel, with a g-factor of 1.

**Figure 2.**
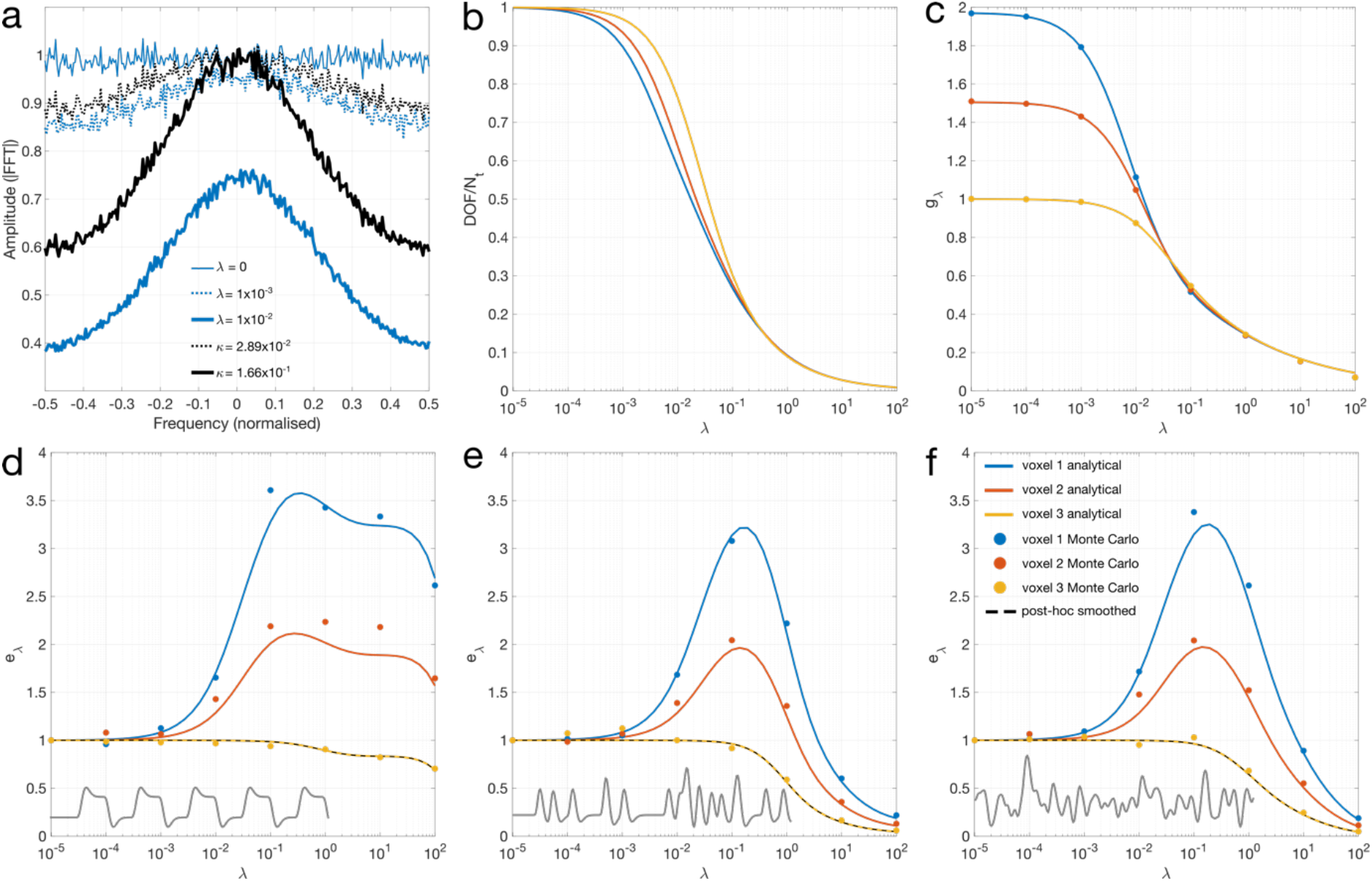
(a) Average spectra from a representative voxel (voxel 1) showing unregularized (thin blue), λ = 1 × 10^−3^ (dotted blue), and *λ* = 1 × 10^−2^ (thick blue) reconstructions. DOF-matching post-hoc kernel smoothing reconstructions are shown in black dotted and thick lines respectively. (b-f) Analytical plots for 3 voxels showing different behavior (blue, orange, yellow lines) are shown alongside corresponding Monte Carlo estimates (blue, orange, yellow dots), versus *λ* (b) Approximate *DOF/N_t_* plots. (c) g-factor. (d-f) Exact GLM efficiency combining information from DOF and g-factors, calculated using regressors (shown in the bottom left) corresponding to (d) block design, (e) event-related design and (f) white noise respectively. DOF-matched post-hoc smoothing efficiency is plotted in dashed black, overlapping with voxel 3.

The relative efficiency metrics (Eq. 9) in Figs. 2d-f, which reflect the ratio of the squared g-factors to the DOF, show that while a net benefit (*e_λ_* > 1) is observed in the linear smoothing reconstructions for any voxel that has a *g*_0_ > 1, there is no gain in efficiency for any voxel that does not alias onto another voxel, where *g*_0_ = 1. Post-hoc smoothing (black dashed line) shows the same efficiency curve as voxel 3, indicating that no efficiency gain is realized. In all cases with an efficiency benefit, *e_λ_* increases with *λ*, to a maximum in this case near *λ* = 10^−1^. As *λ* gets too large, the loss of DOF in the regression begins to outweigh the reduced g-factors, and statistical power decreases. The efficiency curves for each regressor are unique due to the dependence of DOF on the design matrix. However, they all show very similar characteristics for low values of *λ*. They exhibit fairly broad peaks in a logarithmic scale, indicating that near-optimal gains are easily achieved within an order of magnitude of the maxima.

The single-slice digital phantom (*Sim2*) is shown in Fig. 3a. To assess residual leakage and aliasing, the normalized root mean square error (NRMSE) across time was computed across varying *λ* in Fig. 3b, by comparing the reconstructed output to the input signal. While an unbiased reconstruction (*λ* = 0) is zero everywhere, with increasing *λ*, increased intra-slice error and slice leakage is observed. The 95^th^ percentiles for the NRMSE values in each case were: (*λ* = 0, *NRMSE* = 0), (1 × 10^−3^,0.006), (5 × 10^−3^,0.020), and (1 × 10^−2^,0.029) respectively. In Figs. 3c-e, g-factors, *DOF/N_t_*, and efficiency are shown. The voxel-wise maps show that the greatest efficiency increases are in regions with the highest unregularized g-factors, which can be seen when comparing row 1 of Fig. 3c with row 4 of Fig. 3e. Fig. 3f plots a representative central voxel time-series from the digital phantom showing increasing temporal smoothness in the time-courses as *λ* increases. In contrast, Fig. 3g plots time-series from a representative peripheral voxel, which shows very little apparent smoothing.

**Figure 3.**
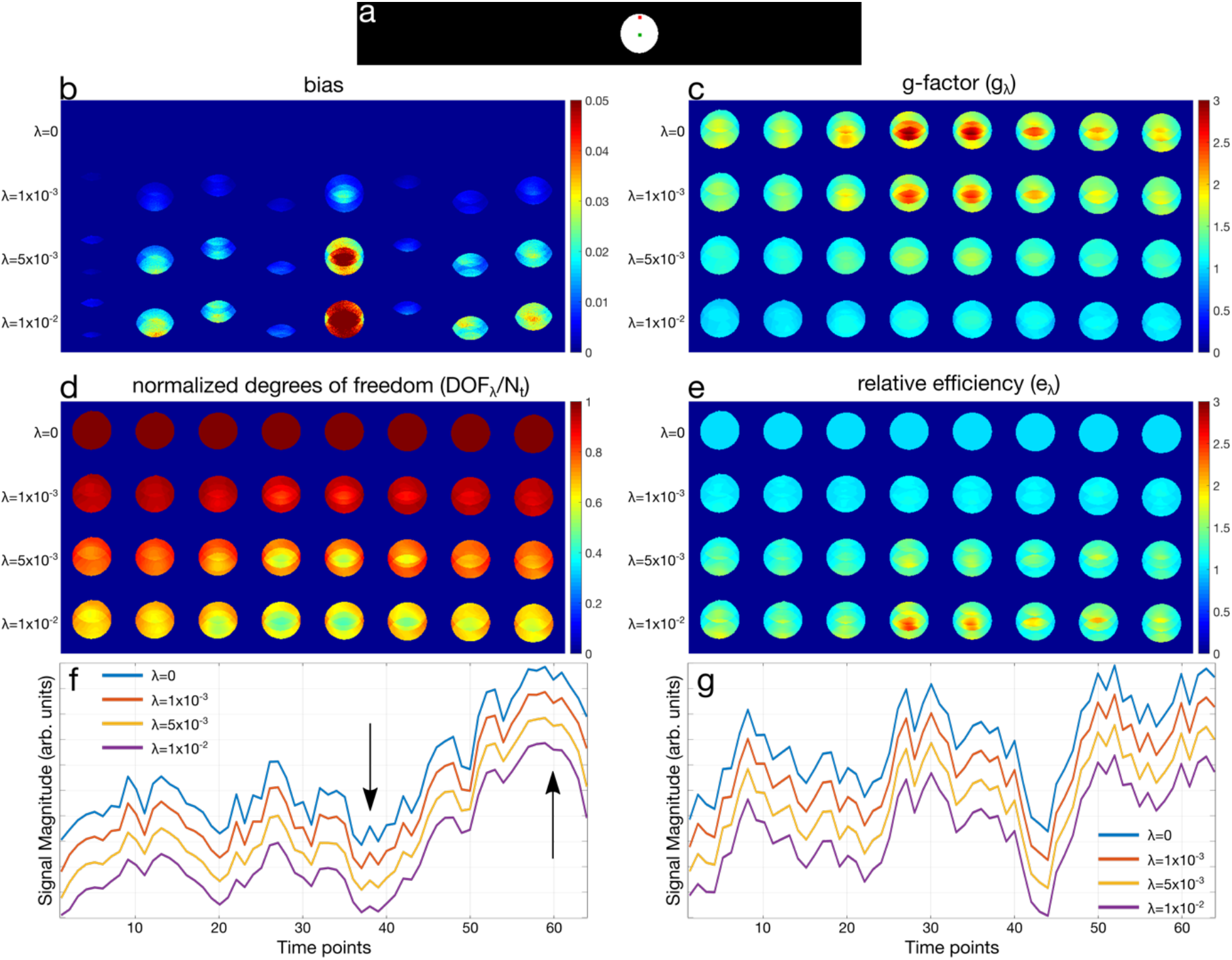
(a) Single slice phantom signal simulation (*Sim2*). Green and red markers denote central and peripheral voxels used in (f,g). (b) Normalized root-mean-square error across time, quantifying spatial leakage bias, with (c) g-factor (d) normalized degrees of freedom (*DOF/N_t_*) (e) GLM relative efficiency. (f,g) Example voxel time-courses for a central and peripheral voxel respectively. The *DOF/N_t_* values are 1.00, 0.81, 0.54, 0.45 as *λ* increases from 0 to 10^−2^ in the central voxel, whereas the *DOF/N_t_* values are 1.00, 0.97, 0.86, 0.76 for the peripheral voxel. Arrows in (f) denote regions where temporal smoothing is evident.

In Fig. 4, the effect of different sampling patterns (at *λ* = 5 × 10^−3^) is shown against fixed, time-independent sampling Δ*k_z_*/Δ*t* = 0, and sampling with shifts of 1, 2, or 3 in the *k_z_* direction. Fig. 4a shows that while residual bias and slice leakage is strongly dependent on *λ*, it is not noticeably different across different Δ*k_z_*/Δ*t* sampling shifts. This is related to the fact that the aliasing pattern (FOV shift) does not change when the entire sampling pattern is shifted across time (Δ*k_z_*/Δ*t* > 0). A reduction in g-factors (Fig. 4b) with time-varying sampling is shown, without much difference in *DOF/N_t_* (Fig. 4c). The net effect on efficiency (Fig. 4d), however, particularly in the (Δ*k_z_*/Δ*t* = 1, 2) cases, is a result of lower g-factors while retaining the same degrees of freedom. Despite the clear benefit of time-varying sampling, the top row of Fig. 4d shows that there is still a small efficiency gain with time-independent sampling.

**Figure 4.**
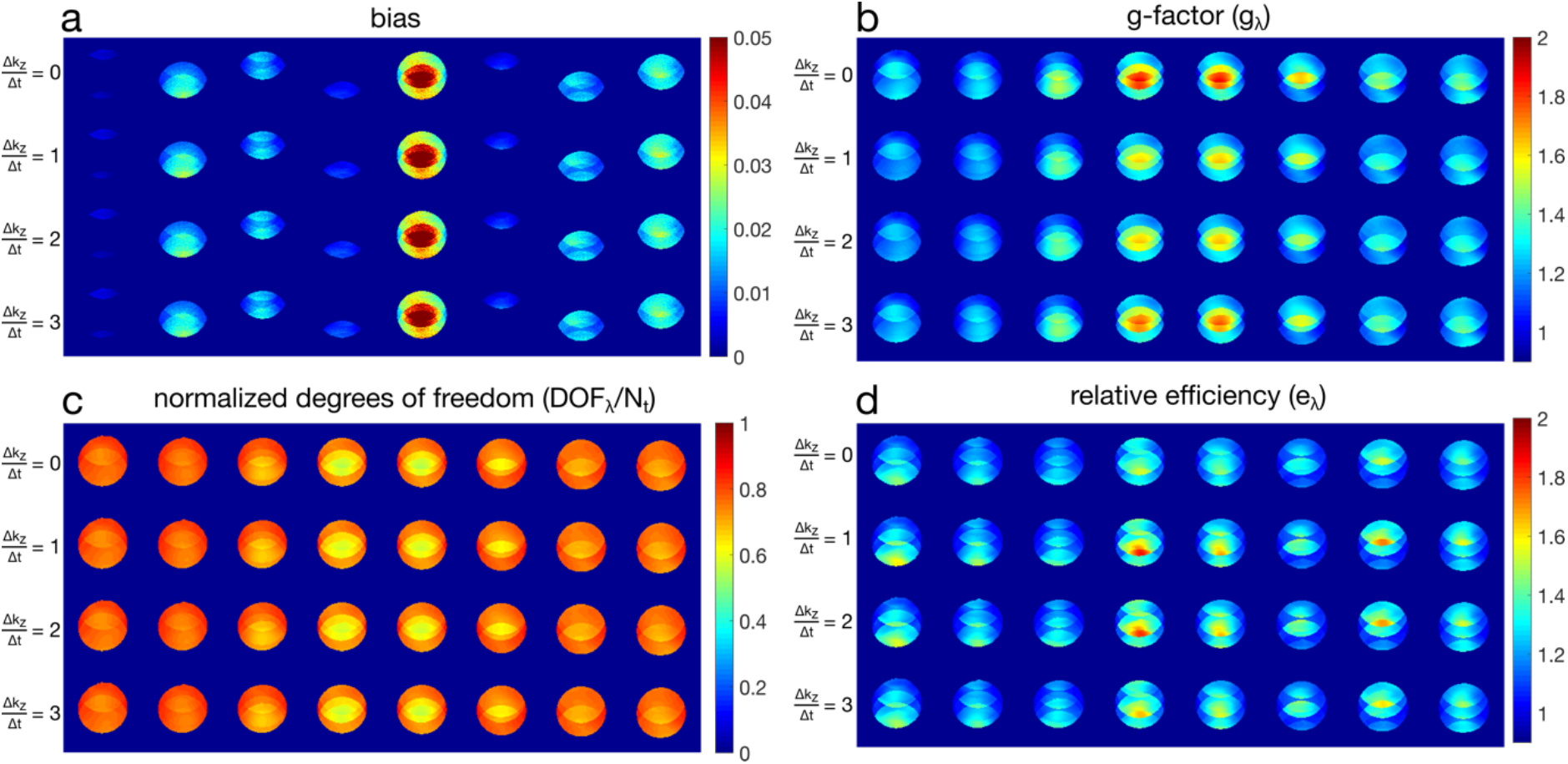
Digital phantom characteristics with different temporal sampling patterns. (a) bias or residual aliasing and slice leakage (b) g-factor, (c) *DOF/N_t_*, (d) GLM relative efficiency.

Single-band excitation *in vivo* data from *Exp1* were reconstructed using the full SMS pipeline for a single subject using the MB8R1 protocol is shown in Fig. 5. Ideally, these data should only signal in the excited slice, although in practice both noise (amplified by g-factors) and slice leakage bias will contribute to the RMSE in the un-excited slices. This data was also reconstructed using conventional single-slice methods (without the slice unaliasing step), to use as a reference for the RMSE calculation, and to normalize the RMSE values. Note that the masking of the coil-sensitivities here produces a brain-like outline in the non-excited slices. The errors are notably smoother in the slice GRAPPA reconstruction outside the target slice, with the regularized reconstructions showing lower overall RMSE, outside of some high RMSE edge or boundary features in the *λ* = 1 × 10^−3^ reconstruction. The errors are visibly noise (g-factor) dominated in the slice GRAPPA (row 1) and *λ* = 1 × 10^−3^ (row 2) reconstructions, and even in the highest regularization reconstruction at *λ* = 1 × 10^−2^ (row 3), little visible aliasing of the excited slice 5 is apparent in any of the other slice locations.

**Figure 5.**
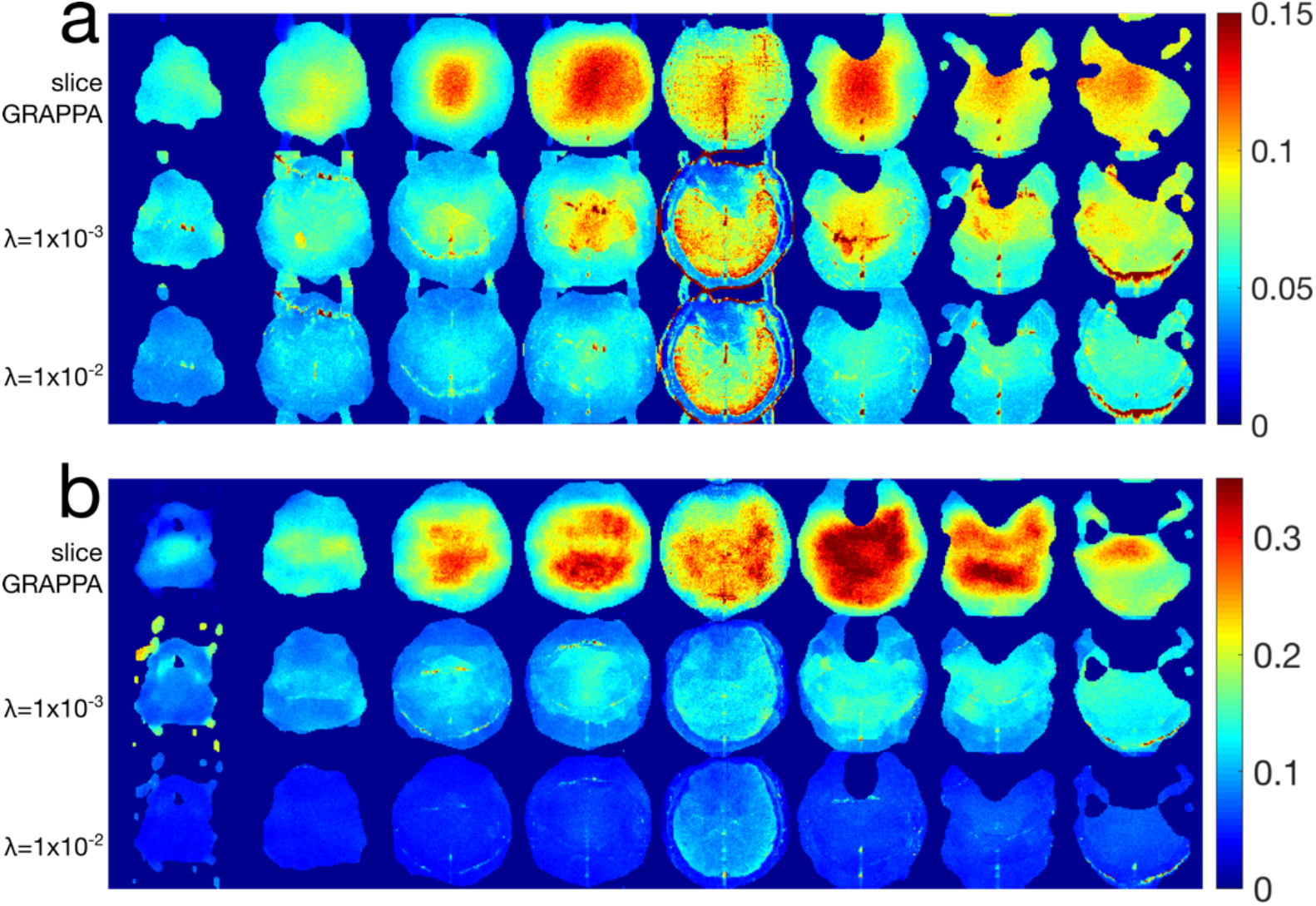
Results from *Exp1* showing the normalized RMSE computed from an SMS reconstruction of singleband excitations of slice 5, using the (a) MB8R1 and (b) MB8R2 acquisition schemes. The RMSE values are normalized to the average RMS signal in the excited slice.

The results of the tSNR efficiency experiments (*Exp2*) are shown in Figures 6 and 7. Fig. 6a shows representative images from the fully excited MB8R1 protocol, across the slice GRAPPA and regularized reconstructions. The tSNR efficiency (Fig. 6b) is slightly improved in central brain areas for the regularized (*λ* = 1 × 10^−2^) reconstruction, which is highlighted by the positive values in the ΔtSNR efficiency maps relative to slice GRAPPA. However, some regions do show reduced tSNR efficiency, particularly noticeable in regions with unaliasing errors, highlighted by the arrows in the example images and tSNR efficiency maps. The arrows denote a region with residual aliasing to incorrect sensitivity estimation of the left eye, corresponding to the most significant region of tSNR efficiency loss.

**Figure 6.**
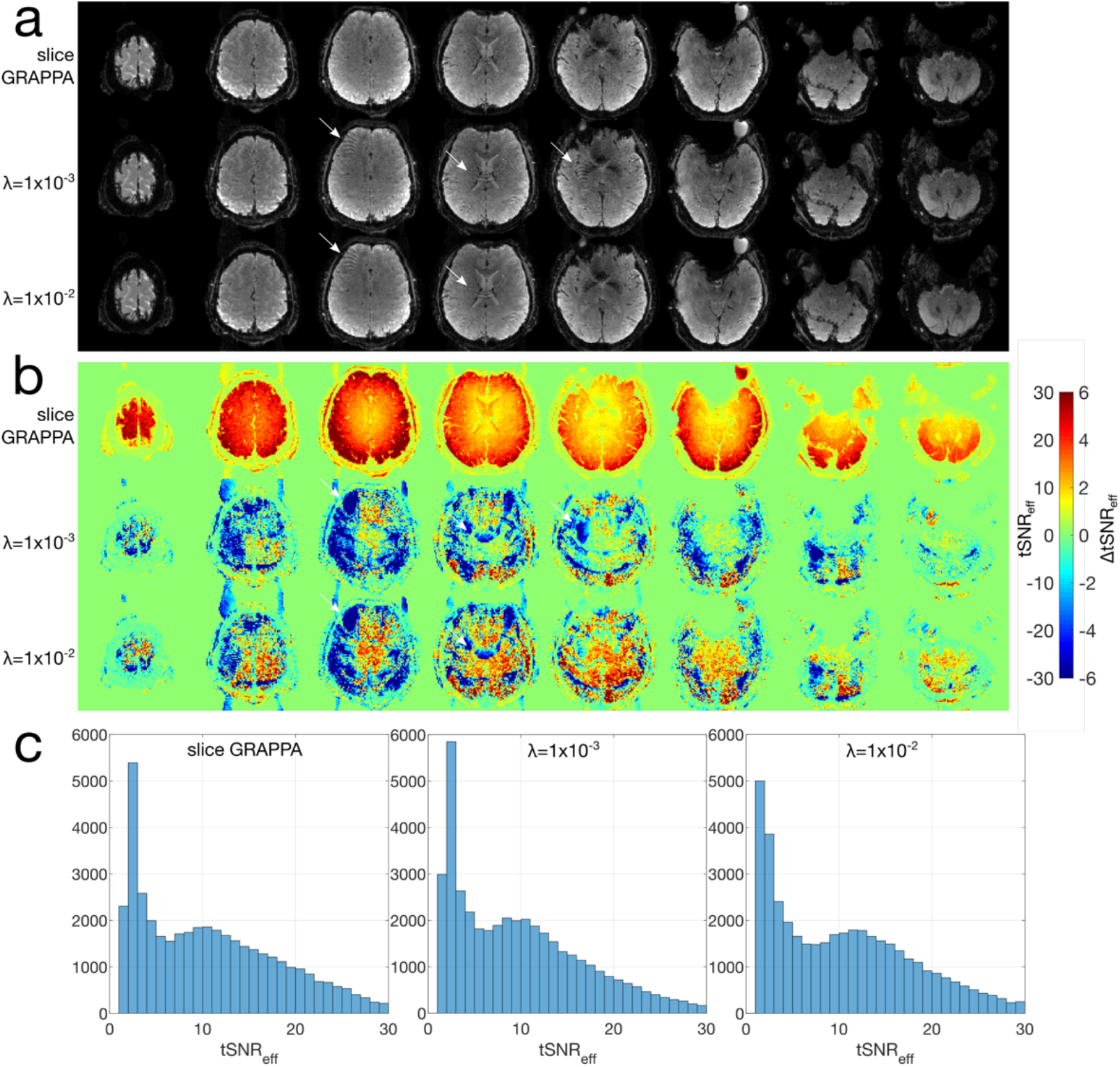
Representative single time-point reconstructed images (a) and tSNR efficiencies (b) for the MB8R1 protocol. The top rows shows the slice GRAPPA reconstruction, followed by the (*λ*= 1× 10^−3^) and (*λ* = 1 × 10^−2^) regularized reconstructions. The tSNR efficiencies in the bottom 2 rows of (b) are shown as ΔtSNR efficiencies relative to slice GRAPPA. The arrows denote regions where unaliasing errors are apparent, and the corresponding regions in the tSNR efficiency maps.

Similar results for the MB8R2 protocol are plotted in Figs. 7a and b, illustrating that the slice GRAPPA reconstruction struggles to robustly unalias the SMS data at this acceleration. However, the temporally regularized reconstructions (*λ* = 1 × 10^−3^,1 × 10^−1^,) result in better images and higher tSNR efficiencies. In both the MB8R1 and MB8R2 regularized reconstructions, sharp boundaries and narrow regions of tSNR efficiency loss can be observed in the tSNR efficiency maps, further highlighting the dependence of the reconstruction on high fidelity sensitivity estimates. To characterize the specific tSNR efficiency benefit in the central brain areas that often suffer most from g-factor penalties, masks were drawn over the deep gray nuclei including the basal ganglia and thalamus (see mask in Supporting Figure S3) for the MB8R2 data. The mean ± standard deviation of the tSNR efficiency for the various reconstructions were, slice GRAPPA: 2.74 ± 0.31, *λ* = 1 × 10^−3^: 4.56 ± 1.03, and *λ* = 1× 10^−2^: 9.02 ± 2.42.

**Figure 7.**
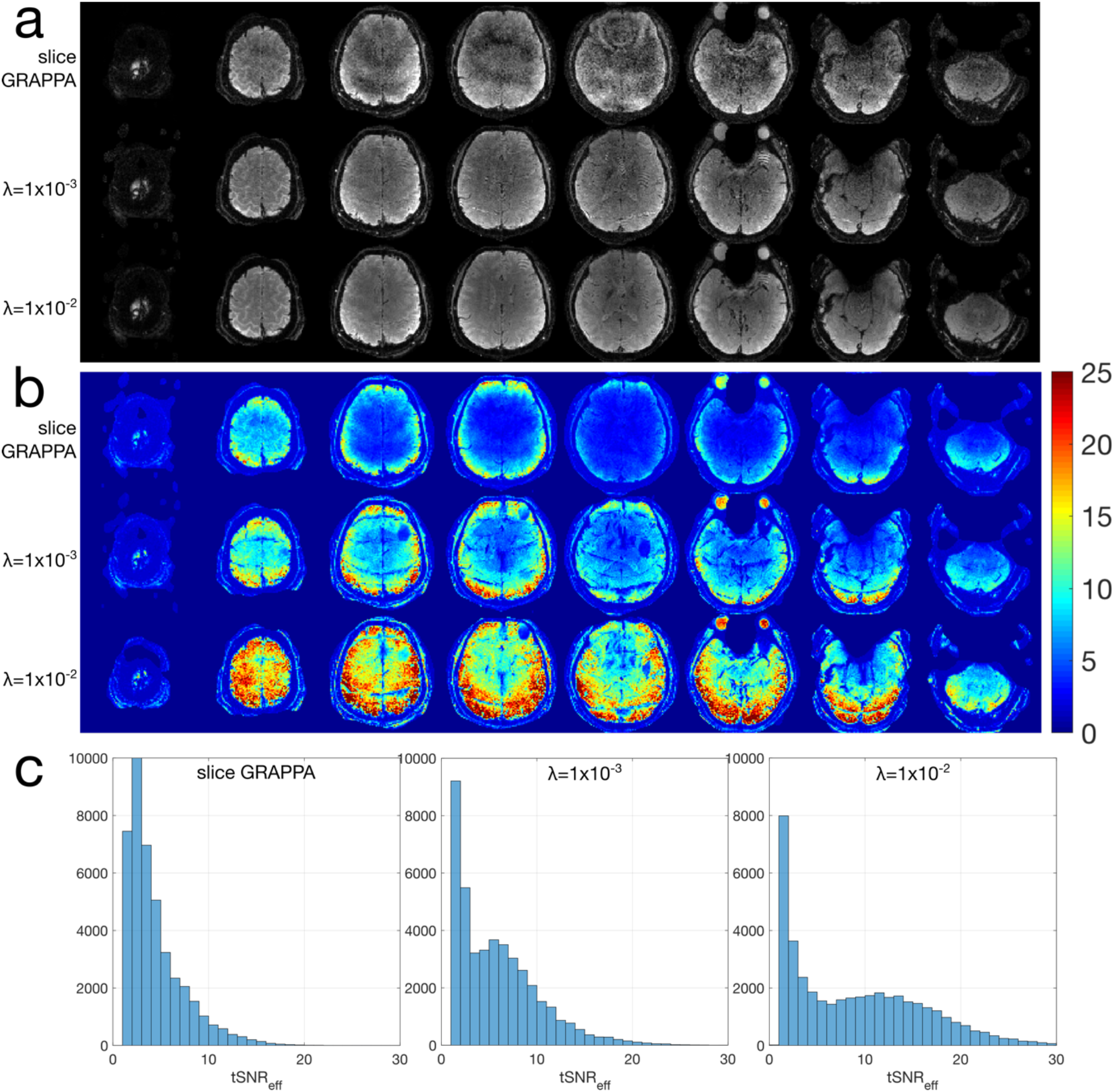
Representative single time-point reconstructed images (a) and tSNR efficiency (b) for the resting MB8R2 protocol. The top row shows the slice GRAPPA reconstruction, followed by the *λ* = 1× 10^−3^ and 1 × 10^−2^ reconstructions. (c) shows histograms of the tSNR efficiency for each dataset.

A comparison of the impact of slower acquisition schemes on tSNR efficiency is shown in Fig. 8, computed analytically using the MB8R2 protocol and the coil sensitivities measured from the subject in Fig. 7. Two “slow” acquisitions were simulated to determine whether the reduction in expected noise variance due to the slower acquisition will compensate for the corresponding reduction in temporal degrees of freedom: (i) an effective MB8R1 acquisition by binning multiple consecutive shots together, and (ii) reducing the number of simultaneous slices down to 4 for a MB4R2 acquisition. Both of these methods halve the effective number of time points (and temporal DOF) in the acquisition, and are compared to the MB8R2 protocol with *λ* = 1× 10^−2^ regularization. To ensure a fair comparison, the MB8R1 efficiency map was multiplied by a factor of 2 to account for the increased sampling, and the MB4R1 efficiency map was increased by a factor of 1.96 to account for higher steady-state signal magnitude with the longer TR (assuming TR = 480 and 960 ms for the MB8R2 and MB4R2 respectively, and T1 = 1600 ms). The results indicate that at comparable effective DOF of approximately 50% relative to the unregularized MB8R2 acquisition in all cases, the *λ* = 1× 10^−2^ MB8R2 reconstruction produces considerably higher efficiency in the central part of the FOV, while performing similarly in the periphery.

**Figure 8.**
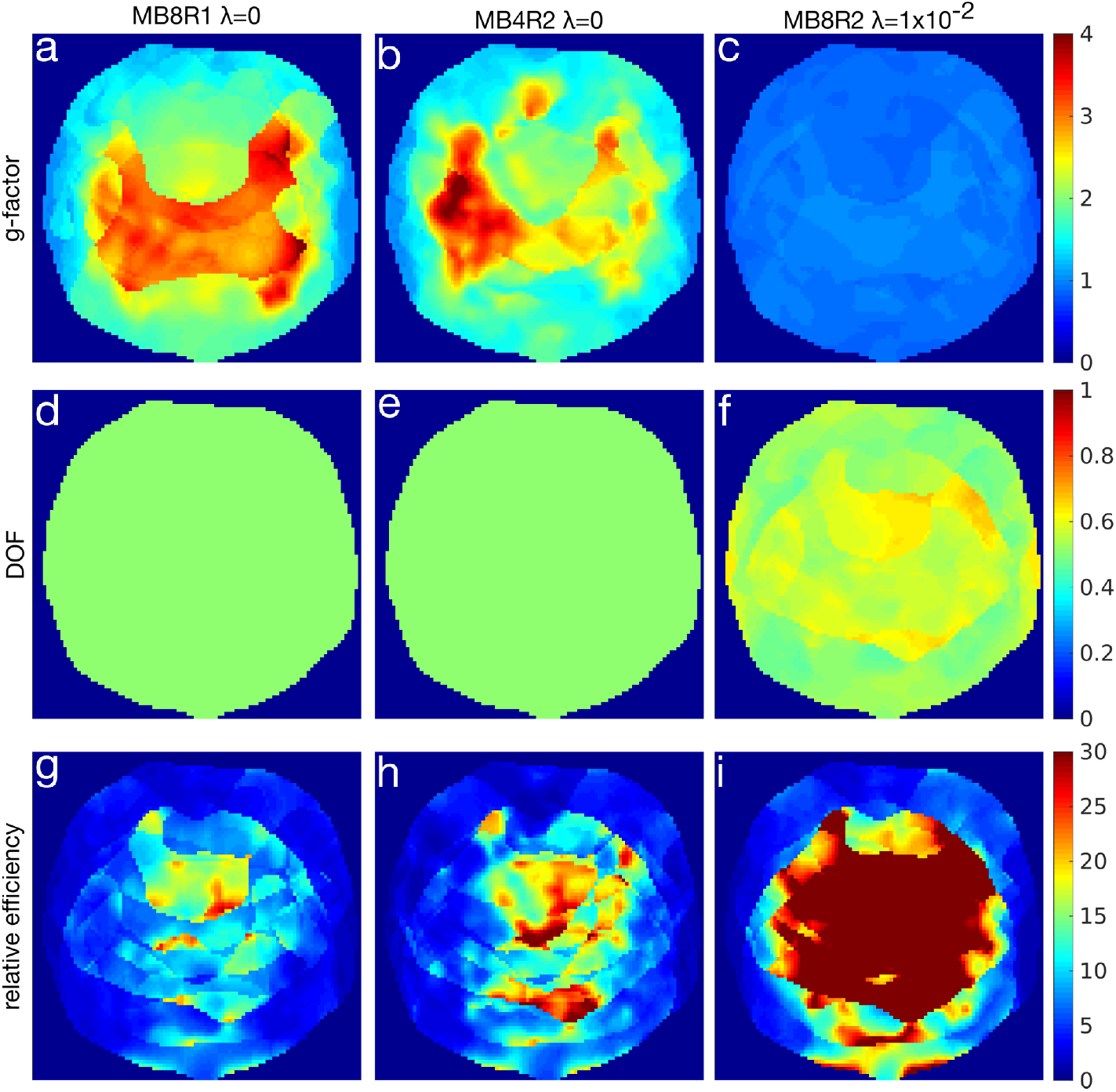
Comparison of g-factors (a-c), normalized DOF (d-f), and relative efficiency (g-i) of 3 different acquisition schemes compared to an unregularized MB8R2 acquisition in a single representative slice. The left column (a,d,g) shows an effective MB8R1 acquisition formed by combining consecutive time-points of the MB8R2 acquisition. The middle column (b,e,h) shows a MB4R2 acquisition that reduces the number of simultaneous slices acquired by half. In both the MB8R1 and MB4R2 cases, the efficiencies have been adjusted to account for the increased sampling or T1 recovery due to the longer volume TR. These are compared to a MB8R2 reconstruction with regularization parameter *λ* = 1 × 10^−2^ in the right column (c,f,i).

The output of GLM analyses on the visual task block-design experiments (*Exp3*) are shown in Figures 9 and 10. In Fig. 9, a MB8R1 dataset is shown for slice GRAPPA and a *λ* = 1 × 10^−2^ regularized reconstructions. At this under-sampling factors, the slice GRAPPA reconstruction is quite robust, and there is little difference in the z-stat maps between the two reconstructions. A scatterplot of the z-statistics in Fig. 9e, however, do reveal a small improvement in z-statistics for the proposed reconstruction over slice GRAPPA, reflecting the increased statistical efficiency.

**Figure 9.**
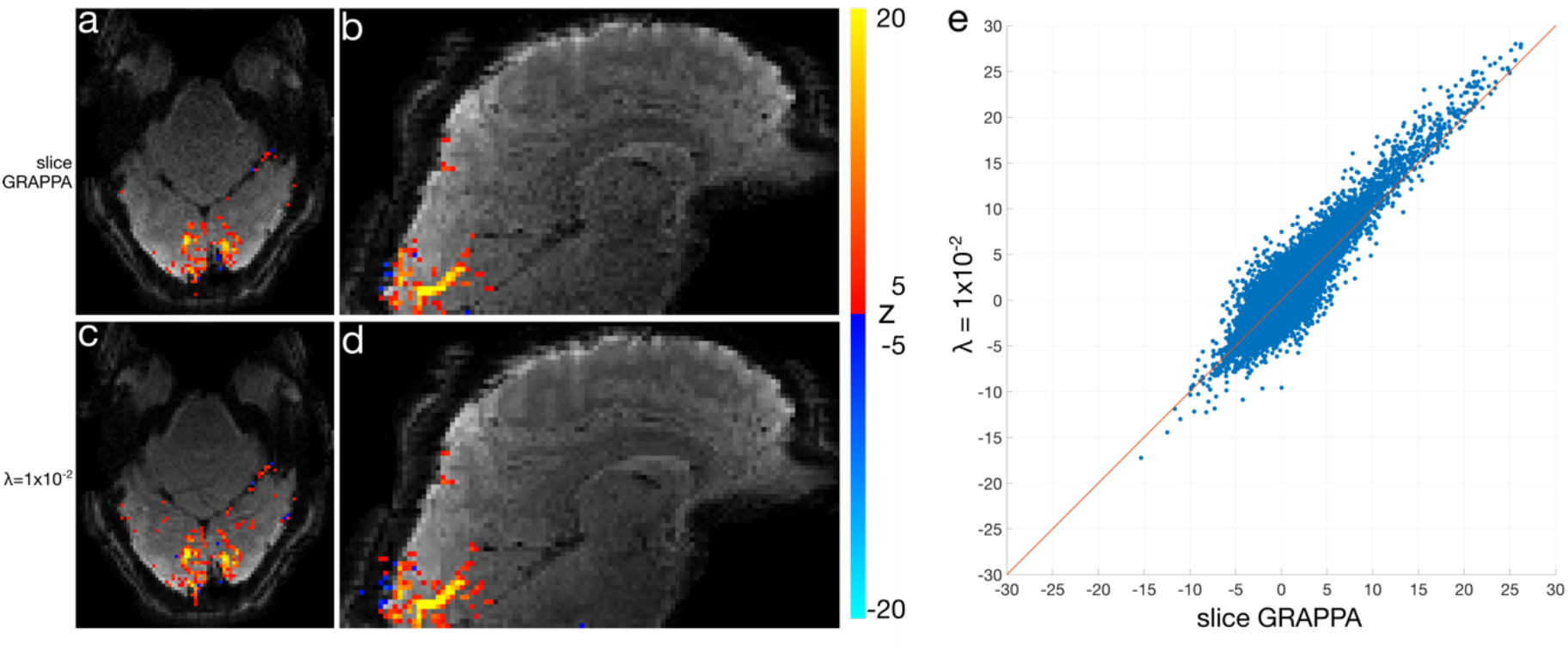
Visual task fMRI dataset using MB8R1 protocol. (a,b) Axial and sagittal z-statistic maps (|*z*| ≥ 5) overlaid on an example image reconstructed using slice GRAPPA. (c,d) Corresponding slices to (a,b) from *λ* = 1 × 10^−2^ regularized SENSE reconstruction. (e) z-statistic scatterplot of all voxels in the *λ* = 1 × 10^−2^ reconstruction against the slice GRAPPA, with the line of identity in orange. Points above the line indicate higher z-statistics in the *λ* = 1× 10^−2^ reconstruction compared to slice GRAPPA.

In Fig. 10, a MB8R2 dataset is shown for slice GRAPPA, *λ* = 1 × 10^−3^, *λ* = 1 × 10^−2^, and post-hoc smoothing reconstructions. Here, the underlying images highlight the apparent tSNR difference expected between the slice GRAPPA and regularized reconstructions, and no residual aliasing is apparent in the images or in the z-statistic maps in any of the reconstructions. While apparent false positives are similar across reconstructions, the regularized reconstructions show much higher sensitivity in the expected primary visual areas. The post-hoc smoothing used a kernel width parameter (*κ* = 2.5 × 10^−1^) chosen to match the average DOF from the *λ* = 1 × 10^−2^ reconstruction. Although the underlying image does appear less noisy, as predicted, no apparent statistical benefit is observed with post-hoc smoothing because the variance per DOF is unchanged. Fig. 10c shows example time-courses from a high z-stat voxel across reconstructions, showing the effect of the reconstruction parameters on temporal fidelity. In Fig. 10d, the power spectra corresponding to these time-courses are shown, with the relatively flat slice GRAPPA spectrum highlighting the fact that these data are dominated by g-factor noise amplification. Furthermore, we see the effect of smoothing as a coloring of the spectra, along with some temporal artefacts (red circles) resulting from the time-varying sampling.

**Figure 10.**
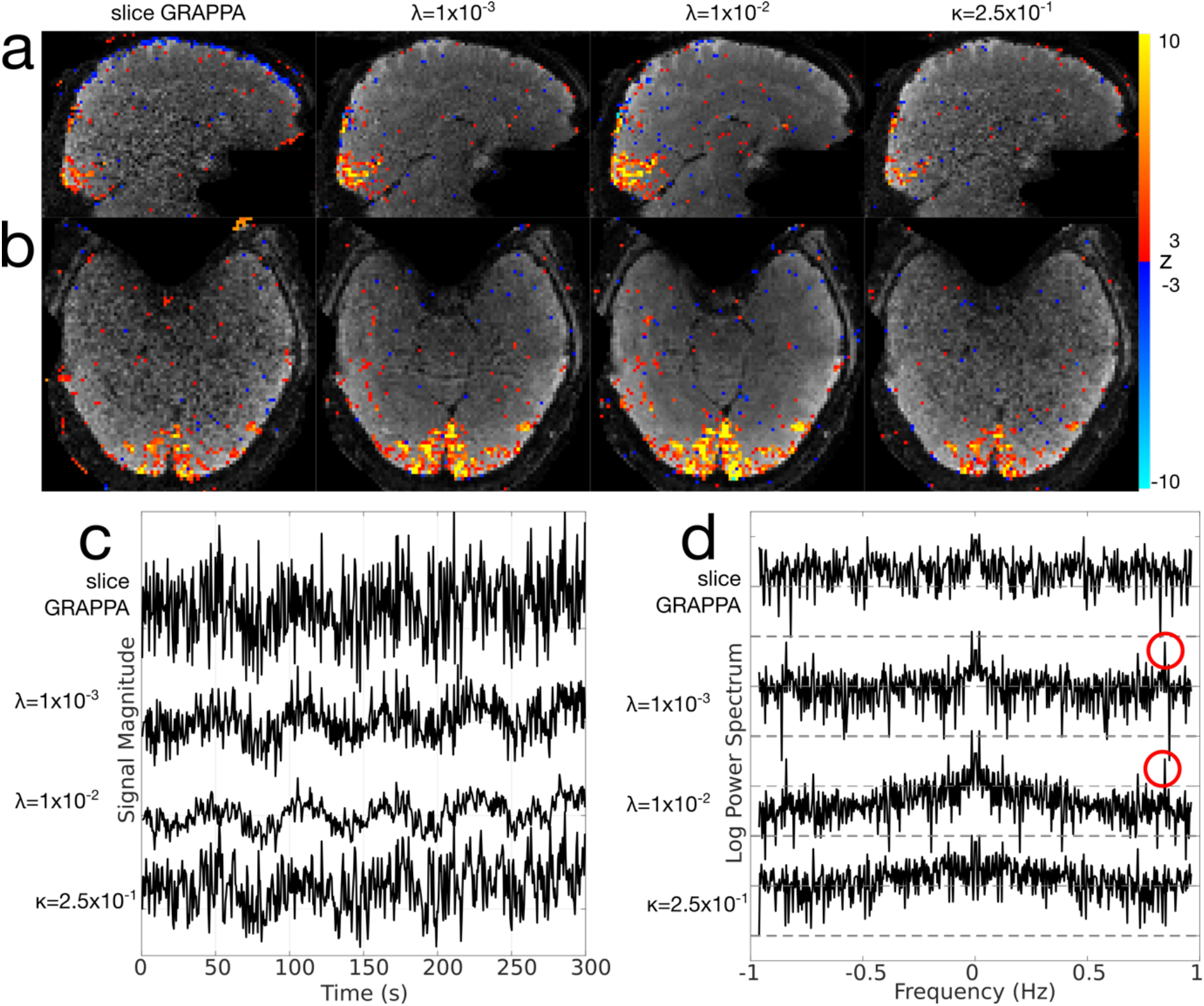
Visual task fMRI dataset using the MB8R2 protocol. (a,b) Sagittal and axial z-statistic maps (|*z*| ≥ 3) overlaid on example image reconstructed images. From left to right, each column represents slice GRAPPA, *λ* = 1× 10^−3^ and *λ*= 1× 10^−2^ regularized reconstructions, and *κ* = 2.5 × 10^−1^ post-hoc smoothed reconstructions. (c) Example time-courses from a single high z-stat voxel, across all reconstructions. (d) Example log power spectra from the same voxel, highlighting the noise colouring resulting from the regularization and post-hoc smoothing. Red circles denote spikes related to the residual bias and the spatio-temporal point spread function.

Results from *Exp4* are shown in Figure 11, highlighting both reconstructed magnitude images at a representative time-point, and tSNR efficiency maps. The results of the slice GRAPPA and *λ* = 1 × 10^−2^ reconstructions are comparable in the “MB8 – 50% gap” protocol, consistent with the MB8R1 results from the previous experiments, even at a reduced slice thickness. However, in the “MB12 – no gap” protocol, the increased reconstruction burden leads to considerable noise amplification in the slice GRAPPA acquisition, which is improved in the *λ* = 1 × 10^−2^ reconstruction, resulting in higher tSNR efficiency, particularly in central brain and cerebellar regions.

**Figure 11.**
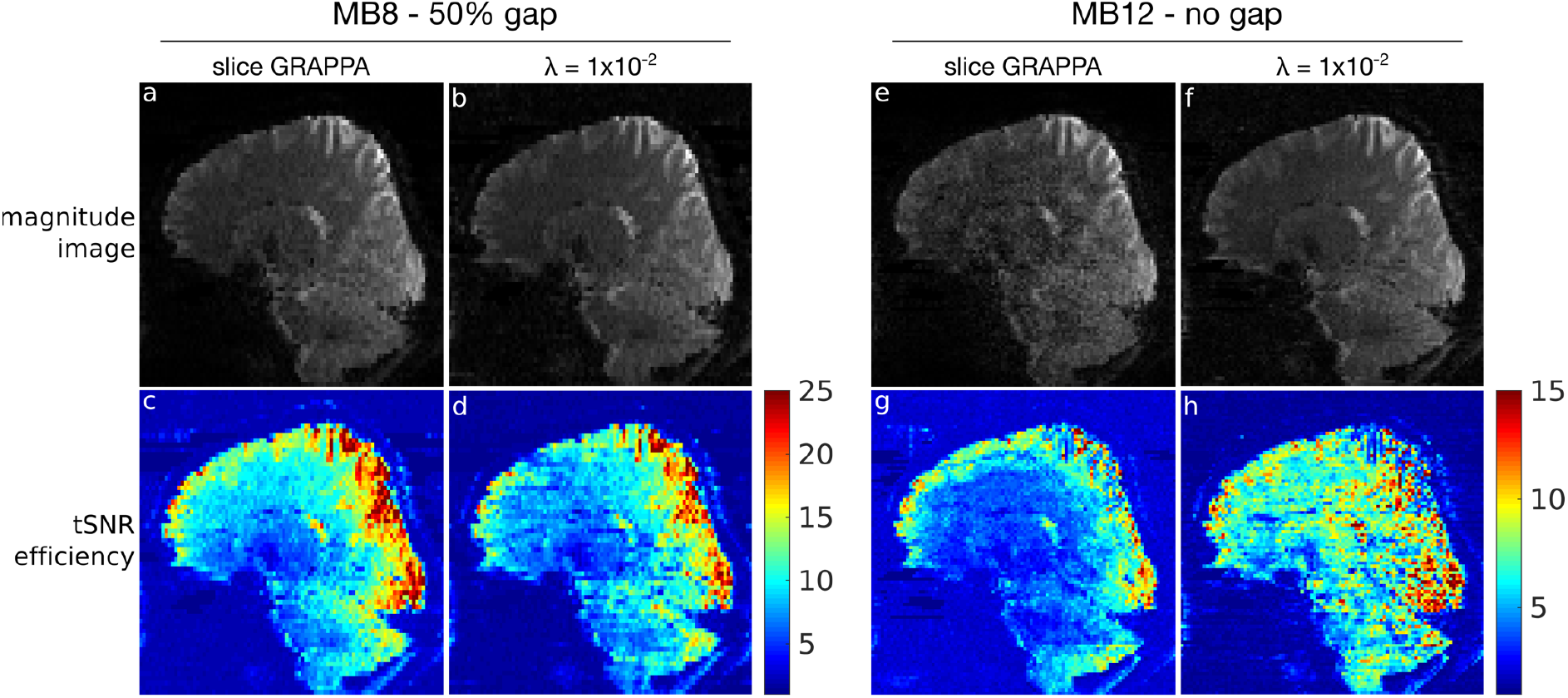
Results of *Exp4*. (a-d) Data from the “MB8 – 50% gap” protocol, and (e-h) data from the “MB12 – no gap” protocol. Within each dataset, the slice GRAPPA reconstruction is shown on the left column, and the regularized reconstruction with *λ* = 1 × 10^−2^ on the right. The top row (a,b,e,f) shows example magnitude images at a single reconstructed time-point, indicating the full-brain and cerebellar coverage. The bottom row (c,d,g,h) shows the tSNR efficiency maps – note the colour scale difference in (c,d) compared to (g,h).

## 5. Discussion

Reconstruction of SMS fMRI data has predominantly been performed using time independent, volume-by-volume reconstructions based on SENSE or GRAPPA-based equivalents. Here, we propose the use of regularization that modulates temporal smoothness in a spatially adaptive manner, to reduce variance (g-factor) by sacrificing temporal degrees of freedom in a way that has a net statistical benefit. Unregularized SENSE reconstructions are unbiased (no leakage) and contrast independent, contingent on the fidelity of the sensitivity estimates. Slice GRAPPA approaches, with limited kernel sizes and dependence on signal contrast changes, trade robustness for residual intra- and inter-slice artefact, effectively allowing for small amounts of bias in the reconstructions to produce better overall images with potentially lower mean squared error. The proposed reconstruction makes a similar trade-off, relying on the low-frequency nature of BOLD signals to ensure that temporal regularization results in comparable or less mean squared error than is produced by conventional slice-GRAPPA unaliasing, while improving statistical efficiency.

One advantage of GRAPPA-based approaches, however, is that explicit estimation of coil sensitivities is not required which makes them more robust to any inconsistencies in the calibration data. In contrast, SENSE based approaches to slice unaliasing depend more strongly on the quality of the estimated sensitivity maps (Zahneisen et al., 2014). Additionally, as can be seen in the tSNR efficiency results, slice GRAPPA produces smoother g-factor noise amplification and therefore smoother tSNR efficiency maps, whereas sharp boundaries or discontinuities can be seen in the proposed SENSE-based reconstructions. These boundaries are due in part to the sharp boundaries in the g-factor and effective DOF maps, characteristic of SENSE reconstructions. Sensitivity mis-estimation, or motion-induced effects, particularly near head boundaries or in regions such as the eyes, can also contribute to localized discontinuities in the tSNR efficiency maps. Increased regularization factors in the proposed method reduce these apparent discontinuities, as the temporal smoothing constraint contributes more to the reconstruction than the sensitivity-encoded data consistency term.

A natural question is whether there is any advantage to acquiring data at higher acceleration factors and temporally smoothing in reconstruction compared to acquisition at intrinsically lower temporal resolution. As we have shown, choosing to reduce DOF by the adaptive smoothing reconstruction does result in higher efficiency than slower acquisition strategies, which reduce DOF uniformly across the volume by taking more time to sample each volume. Although the proposed method results in images with spatially varying DOF, voxel-wise variation in efficiency is already commonplace using conventional parallel imaging due to spatially varying g-factors. Other advantages of acquiring data at higher acceleration factors include reduced intra-volume motion, potentially better contrast, and reduced physiological noise aliasing when slower acquisitions do not sufficiently sample cardiac frequencies.

Spatially varying smoothing is also possible using voxel-wise post-hoc kernel smoothing of varying kernel widths. However, we have shown that post-hoc smoothing with an encoding independent kernel can only reduce the efficiency (Liu and Frank, 2004; Smith et al., 2007). The proposed smoothing regularized reconstruction, in contrast, depends on the encoding (i.e. the k-space sampling pattern and coil sensitivities), and reduces noise variance beyond what would be expected from the spectral filtering effect alone. By shifting the noise spectra down, there is a net reduction in variance per degree of freedom, and therefore a benefit to statistical inference. The efficiency measure presented here generalizes the tSNR efficiency commonly used to assess fMRI time-series fidelity, by accounting for the effective degrees of freedom, rather than acquisition time.

The method described in this paper is not specific to SMS-EPI, although we chose to focus on it in the scope of this work due to its popularity for fMRI data acquisition, and the simplicity of the sampling modifications required. The proposed reconstruction framework does not depend on any specific features of SMS acquisition, and will benefit any 2D or 3D acquisition strategy that can support time-varying sampling schemes. Although time-varying sampling is not a requirement of the method, efficiency gains are significantly greater when time-varying sampling is employed. This is analogous to FOV shifting in conventional CAIPI, by optimizing the sampling scheme to with respect to the reconstruction constraints. In this case, time-varying k-space samples provide more information when the smoothness constraint effectively shares sampling information across time. As we have also shown, the sampling pattern does not need to have a fixed shift across time, although regular sampling does result in the highest efficiency gains. With a fixed shift across time, where Δ*kz*/Δ*t* is constant, the g-factors are time-independent. In the case of more general sampling patterns, as with the pseudorandom sampling, the conditioning of the inverse problem associated with any given time-point is dependent on the sampling of its neighbors, resulting in time-dependent g-factors that can be used to assess the impact of noise amplification on individual time-points.

The smoothing parameter *λ* can be selected using a number of different criteria: an upper limit on the amount of spatial leakage given some signal model, minimizing mean-squared error, maximum efficiency, or a lower limit on the retained DOF. The benefit of the latter two choices is that they are signal independent, although the trade-off between bias and variance in the reconstruction does require a signal model to fully characterize. Simulations showed that while the peak efficiency points can occur at relatively high regularization factors (e.g. *λ* = 1 × 10^−1^), smaller regularization factors (e.g. *λ* = 1 × 10^−2^) can still provide considerable efficiency gains without noticeable bias. The spatio-temporal characteristics of the residual bias depend in part on the k-space sampling scheme, and can be modulated by changing the temporal sampling shifts to produce less temporally coherent residual aliasing, although this can cause issues with data analysis due to non-stationary noise resulting from the time-varying g-factors.

While the assumption of low-frequency signal content was used to justify the use of the temporal regularization by reducing the impact of signal leakage, the efficiency gain only describes the effects on the residual white Gaussian thermal noise, and does not account for noise autocorrelations or physiological noise. While consideration of a more comprehensive fMRI noise model would provide more accurate statistical efficiency modelling, e.g. by including spatio-temporal noise characteristics of sub-second TR acquisitions (Bollmann et al., 2018), they would substantially increase noise model complexity. The presence of other noise sources diminishes the impact of the efficiency gains calculated here, although in g-factor limited regimes where the noise is thermally dominated, the assumed thermal noise model is asymptotically correct. Furthermore in *Exp4*, data acquired at higher spatial resolution across slice acceleration factors demonstrated the benefit of the proposed reconstruction under conditions with controlled temporal autocorrelation (fixed volume TR). However, incorporating more sophisticated noise modelling and better informed constraints for the regularized time-series image reconstruction, as well as interactions between regularization factors and acquisition parameters such as volume TR would be a natural extension of this work.

In this work, we focus on statistical efficiency (reducing noise variance per degree of freedom) as the primary metric of reconstruction quality, rather than optimizing directly for minimizing noise or maximizing SNR. We explicitly construct the efficiency metric expression by an end-to-end consideration of the fMRI dataset, from noisy multi-coil k-space to the variance of the GLM estimator and the tSNR efficiency, which is possible due to the linear reconstruction and analysis framework. While we do not directly optimize for other fMRI analysis methods, such as independent component or seed-correlation analysis for resting state fMRI, reduced noise variance per DOF should also be beneficial, although this has not yet been evaluated.

## 6. Conclusion

A linear reconstruction method for SMS-EPI fMRI data has been developed based on temporal smoothing regularization in a spatially adaptive, encoding-dependent way. This reduces noise for improved statistical power and efficiency compared to conventional slice-GRAPPA reconstructions.

## Appendix A

The reconstruction is a linear transform of the measured k-space and a noise term:

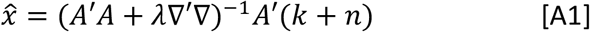

The noise model we assume is zero-mean, Gaussian white noise, with *n* = *N*(0, *σ*), and we further assume that the noise has been whitened using the coil noise covariance, so that:

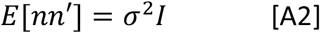

The mean or expectation of the estimate (because the noise is zero mean):

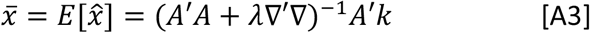

and the mean-subtracted estimate is a function of the noise:

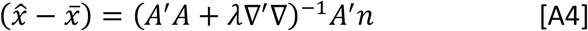

We calculate the g-factor from the voxel-wise variance of the linear estimator:

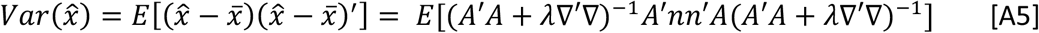

Since the noise term is the only stochastic part of this expression:

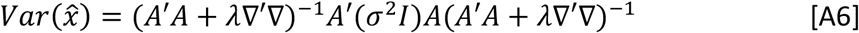

The square root of the entries along the diagonal of 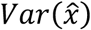 reflect the standard deviation of the noise expected in each voxel. The g-factor is calculated as the ratio of this standard deviation with the standard deviation expected from a fully-sampled acquisition. Since the fully-sampled, un-regularized variance in each voxel *m* is *σ*^2^/(*A’A*)_*m,t*_, the ratio then becomes:

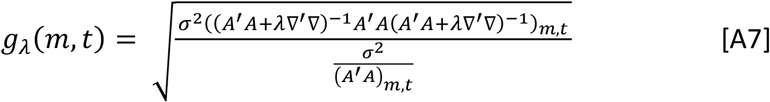

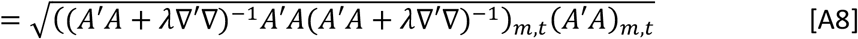

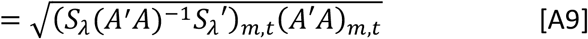

where *S_λ_* = (*A’A* + *λ*∇’∇)^−1^*A’A*, and *X_m,t_* denotes the diagonal element of *X* corresponding to the *m^th^* voxel and *t^th^* time-point.

For the operator ∇’∇: if the data matrix *X* were vectorized with each voxel’s time-course vertically concatenated, ∇’∇ would be a block diagonal matrix where each block is a matrix with mostly 2 on the diagonals, and –1 on the neighboring off-diagonals. An example ∇’∇ diagonal block for a 5-time point input is given as:

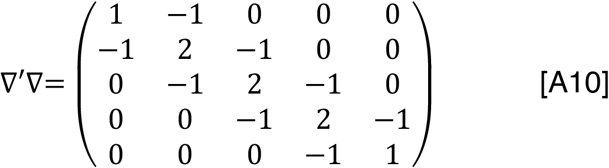

We define ∇ using non-circular boundary conditions (i.e. only finite differences between neighboring points were taken, not including the difference between the first and last time-points). In practice, this matrix was not constructed, but the action of ∇’∇ on *X* was implemented by subtracting temporally shifted versions of *X* to a scaled version of itself, using MATLAB’s circshift function (and taking care of the boundary conditions appropriately).

## Appendix B

In the context of a univariate GLM analysis, a regression coefficient 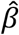 can be expressed as a function of *S_λ_* as:

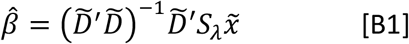

where 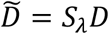 is the smoothed design matrix, assumed to be a single column. More complex contrasts can be accommodated without any loss of generality by a simple transformation (Smith et al., 2007).

From Worsley and Friston (Worsley and Friston, 1995), the variance of the regression coefficient 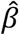 is:

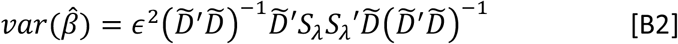

with

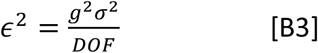

and

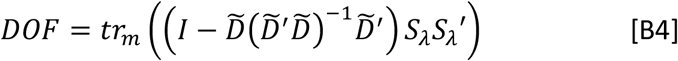

with *tr_m_*(·) used to indicate that values across time for a given voxel *m* are summed. In Eq. [B3], we make use of the fact that following reconstruction, the noise variance is amplified by the square of the g-factor. Note that in Worsley et al., the design matrix and smoothing operators are denoted by *G* and *K* respectively.

Then benefit of regularization, in the context of the GLM, can be quantified by the relative efficiency of the estimator, where a more efficient estimator has increased sensitivity and statistical power. It is defined by the ratio of the variances of the estimators in the unregularized (*λ* = 0) and regularized cases:

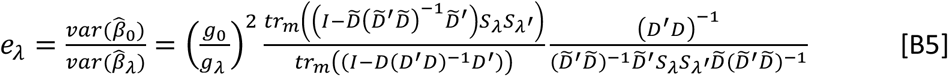

which dictates whether the decrease in noise amplification outpaces the loss in effective DOF. When *e_λ_* > 1, a net benefit in GLM efficiency is observed.

## Acknowledgements

The authors would like to thank Steen Moeller for providing the slice GRAPPA reconstruction code. This research was funded by the Royal Academy of Engineering (MC, RF201617\16\23) and the Wellcome Trust (KLM, 202788/Z/16/Z). The Wellcome Centre for Integrative Neuroimaging is supported by core funding from the Wellcome Trust (203139/Z/16/Z).

