## Supporting Information for "Improved Statistical Efficiency of Simultaneous Multi-Slice fMRI by Reconstruction with Spatially Adaptive Temporal Smoothing"

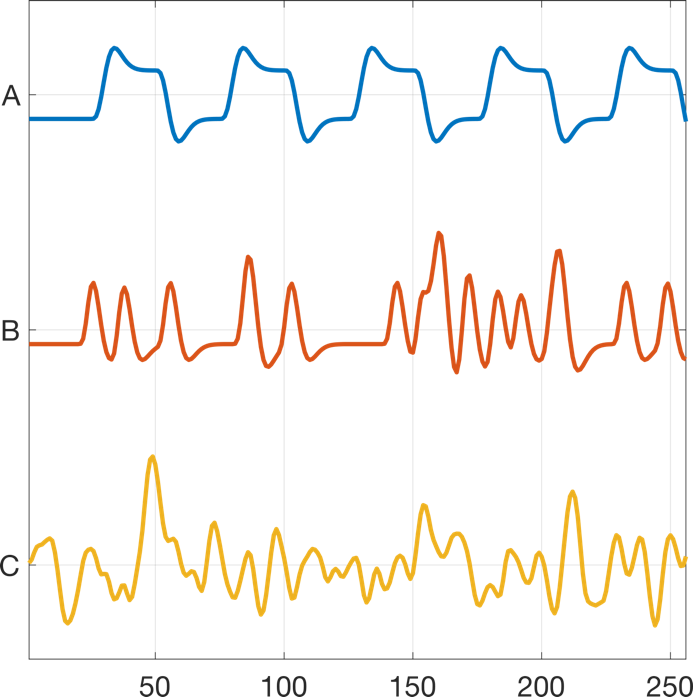

Figure S1 - Example regressors used to assess relative GLM efficiency: (A) 5-period block design task, (B) jittered event-related task, (C) Gaussian white noise regressor. All 3 regressors were convolved with a canonical hemodynamic response function.

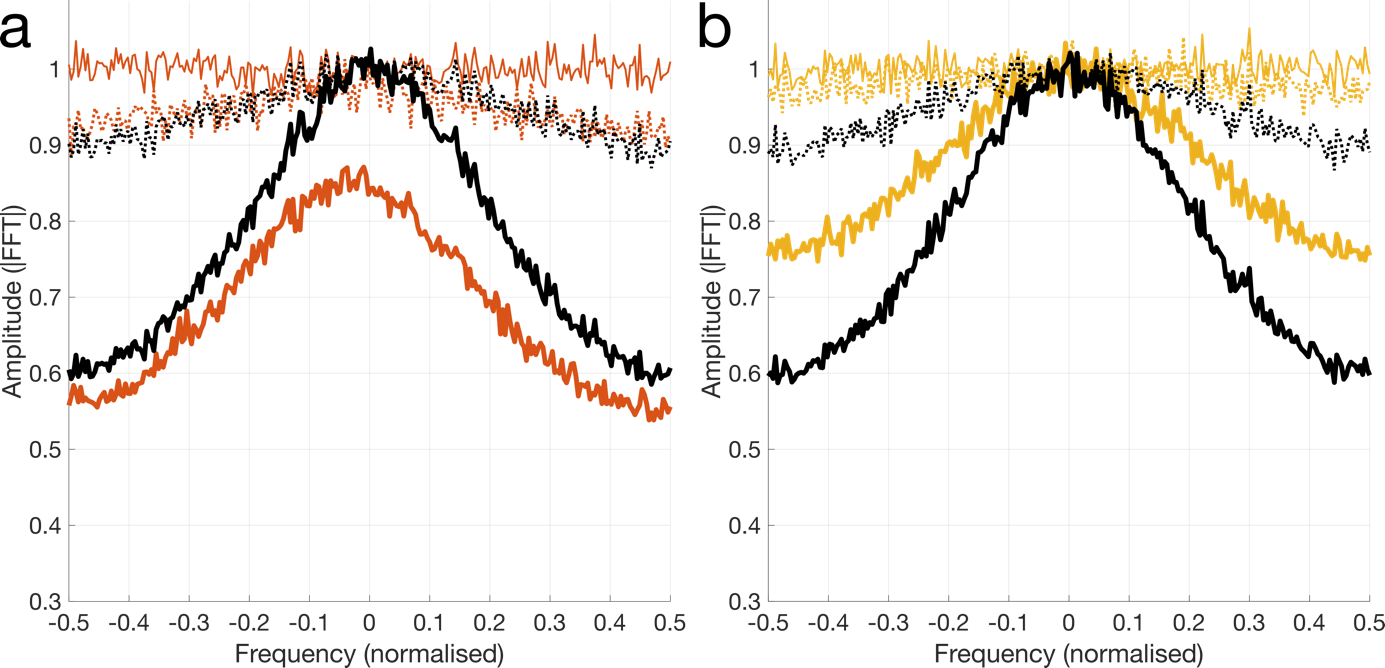

Figure S2 - Noise spectra plotted for additional 2 voxels as a function of of the regularization parameter $\lambda$. Note that the black curves, representing post-hoc kernel smoothing do not change from voxel-to-voxel, and was chosen here to match DOF with the voxel shown in Fig. 2a. Plot color-scheme is matched to those in Fig. 2.

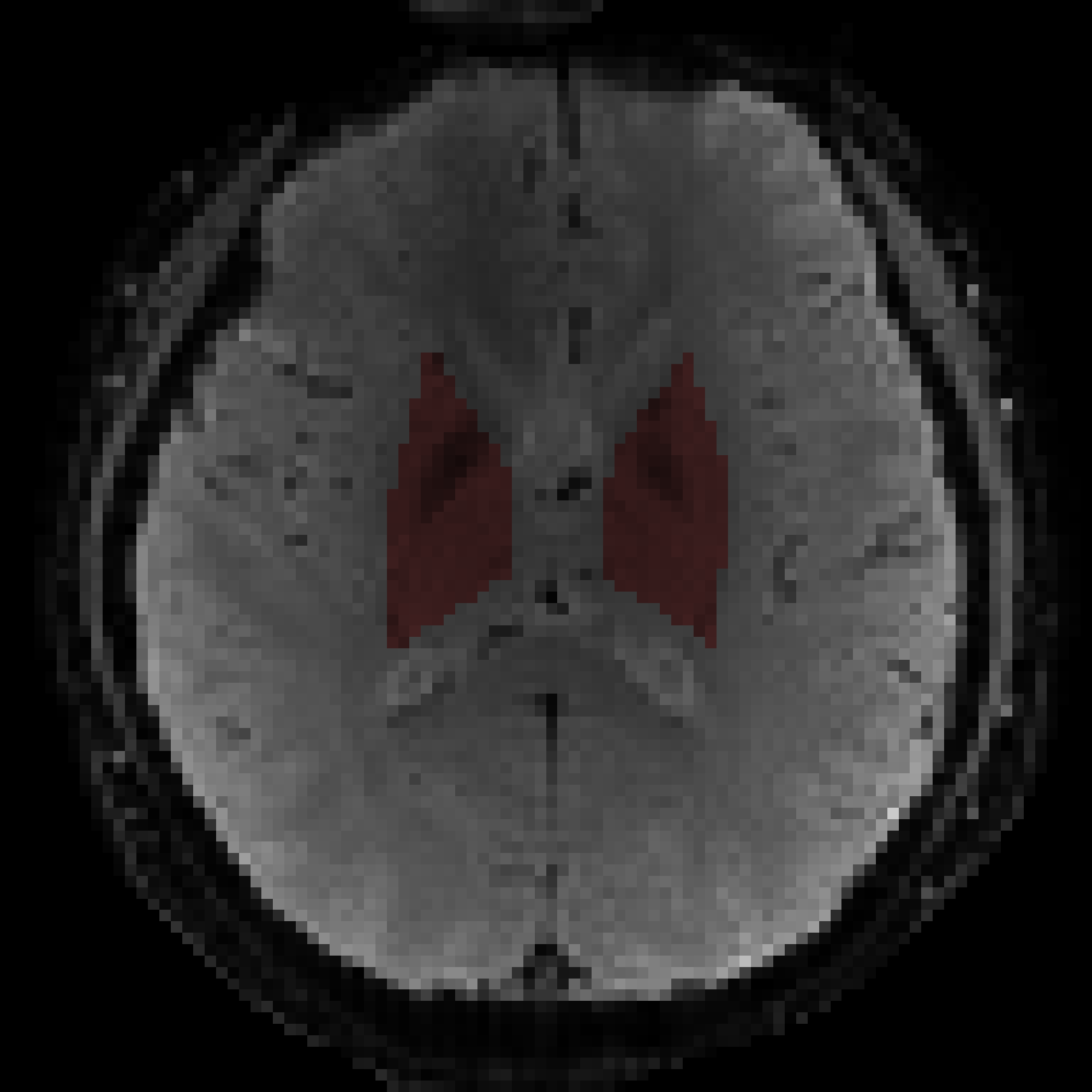

Figure S3 - Deep gray nuceli mask (highlighted in red) including basal ganglia and thalamus, for evaluation of tSNR efficiencies in the MB8R2 data.
